# mTOR signaling controls the formation of smooth muscle cell-derived intimal fibroblasts during vasculitis

**DOI:** 10.1101/2023.12.15.571811

**Authors:** Angus. T. Stock, Sarah Parsons, Jacinta. A. Hansen, Damian. B. D’Silva, Graham Starkey, Aly Fayed, Xin Yi Lim, Rohit D’Costa, Claire. L. Gordon, Ian. P. Wicks

**Affiliations:** WEHI, Melbourne, 3052, Australia; Department of Forensic Medicine, Monash University, Melbourne, 3006, Australia; Victorian Institute of Forensic Medicine, Melbourne, 3006, Australia; Liver & Intestinal Transplant Unit, Austin Health, Melbourne, 3084, Australia; Department of Surgery, The University of Melbourne, Austin Health, Melbourne, 3084, Australia; Department of Surgery, Austin Health, Melbourne, 3084, Australia; Department of Infectious Diseases, Austin Health, Melbourne, 3084, Australia; DonateLife Victoria, Carlton, 3053, Australia; Department of Intensive Care Medicine, Melbourne Health, Melbourne, 3084, Australia; Department of Microbiology and Immunology, The University of Melbourne, The Peter Doherty Institute for Infection and Immunity, Melbourne, 3052, Australia; North Eastern Public Health Unit, Austin Health, Melbourne 3084, Australia; Rheumatology Unit, The Royal Melbourne Hospital, 3050, Australia; University of Melbourne, Department of Medical Biology, 3052, Australia

**Keywords:** Vasculitis, stenosis, fibroblasts, mTOR, Kawasaki Disease

## Abstract

The excessive accumulation of fibroblasts within the intimal layer of inflamed vessels is a feared complication of vasculitis, which can lead to arterial stenosis and ischemia. In this study, we have investigated how such intimal fibroblasts develop during Kawasaki Disease (KD), a paediatric vasculitis typically involving the coronary arteries. By performing lineage tracing studies in a murine model of KD, we reveal that vasculitis-induced intimal fibroblasts develop independently of both adventitial fibroblasts and endothelial cells, and instead derive from smooth muscle cells (SMCs). Notably, the emergence of SMC-derived intimal fibroblasts - in both mice and in patients with KD, Takayasu’s arteritis and Giant Cell arteritis - coincided with their activation of the mechanistic target of rapamycin (mTOR) signalling pathway. Moreover, the genetic deletion of mTOR signalling in SMCs abrogated the emergence of intimal fibroblasts, demonstrating that mTOR is an intrinsic and essential regulator of vasculitis-induced, SMC-derived intimal fibroblasts. Collectively these findings provide molecular insight into the pathogenesis of arterial stenosis and identify mTOR as a therapeutic target to prevent adverse vascular remodelling in vasculitis.

## Introduction

Kawasaki Disease (KD) is a medium vessel vasculitis, now recognised as the leading cause of acquired childhood heart disease in developed countries [1, 2]. The major clinical complication for children with KD is the development of coronary artery disease. This manifests in the first few weeks of disease as coronary artery aneurysms, which in most cases resolve [3]. However, patients who develop large or giant coronary aneurysms are at significant risk of subsequent cardiac events [3] caused by the pathological remodelling of the coronary arteries [3–5]. Indeed a majority of these KD patients develop coronary artery stenosis [4], with a significant risk of myocardial infarction due to occlusion or thrombosis [3, 6, 7].

A similar pathological sequence leading to arterial stenosis can occur in other types of vasculitis, such as Takayasu’s arteritis (TAK) and Giant cell arteritis (GCA) [8–11]. Histological analysis of inflamed arteries has revealed that fibroblasts are the dominant population in the intimal layer of occluded arteries in KD and GCA patients [6, 7, 12–15]. Consequently, vasculitis induced arterial stenosis is usually attributed to the dysregulated infiltration and proliferation of pathogenic fibroblasts into and within the intimal layer of inflamed arteries [2, 6].

How intimal fibroblasts develop and function during vasculitis is not well understood. It has been reported that the phenotype of intimal fibroblasts populating inflamed temporal arteries of GCA patients overlaps with their adventitial counterparts [16], suggesting these populations may be interrelated. This possibility is consistent with earlier observations that adventitial fibroblasts can migrate into the neointima after experimental balloon injury of the carotid artery [17, 18], and argues that intimal fibroblasts develop from migratory adventitial fibroblasts [18]. However, the expression of alpha-smooth muscle actin (α-SMA) by intimal fibroblasts [19] has led to the suggestion that intimal fibroblasts develop from smooth muscle cells (SMCs), similar to what is reported during atherosclerosis [20]. Endothelial cells and pericytes have also been shown to differentiate into myofibroblasts during experimental models of cardiac and kidney disease [21–24], raising the possibility that intimal fibroblasts may develop locally, from such intimal resident precursors. Thus, at present, the source of intimal fibroblasts that emerge in vasculitis remains contentious, and may potentially vary between different diseases, tissues and even blood vessel types.

In this study, we sought to define the origin of intimal fibroblasts that emerge during vasculitis and explore the signalling pathways that control their development. To do so, we have used complimentary lineage tracing systems in a mouse model of KD that re-capitulates coronary artery stenosis in human disease. Our findings reveal that intimal fibroblasts develop from SMCs and their formation (in multiple forms of systemic vasculitis) is driven by activation of the mechanistic target of rapamycin (mTOR) signalling pathway.

## Results

### Collagen-expressing fibroblasts populate the coronary artery intima during CAWS induced vasculitis

To investigate the origin of vasculitis-induced intimal fibroblasts, we employed a murine model of Kawasaki Disease (KD), where cardiac vasculitis is induced by the injection the *Candida albicans* water soluble complex (CAWS)[25, 26]. Histological analysis of cardiac sections revealed that CAWS injected mice (analysed at 4-5 weeks post CAWS injection) developed extensive transmural inflammation of the coronary arteries, which was accompanied by profound collagen deposition (as measured by Sirius red staining) within the adventitial and intimal layers (**Fig 1A**). Moreover, by using confocal microscopy, we found that the endothelial (visualised by CD31 expression) and medial (using autofluorescence to visualise the elastic fibres of the media) layers separate during CAWS induced vasculitis, due to significant thickening of the intimal layer (**Fig 1B**). Thus, similar to what is described in human KD [3, 6, 13], CAWS injection triggers intimal hyperplasia of the coronary arteries, characterised by cellular infiltration and fibrosis within the intimal layer.

**Figure 1:**
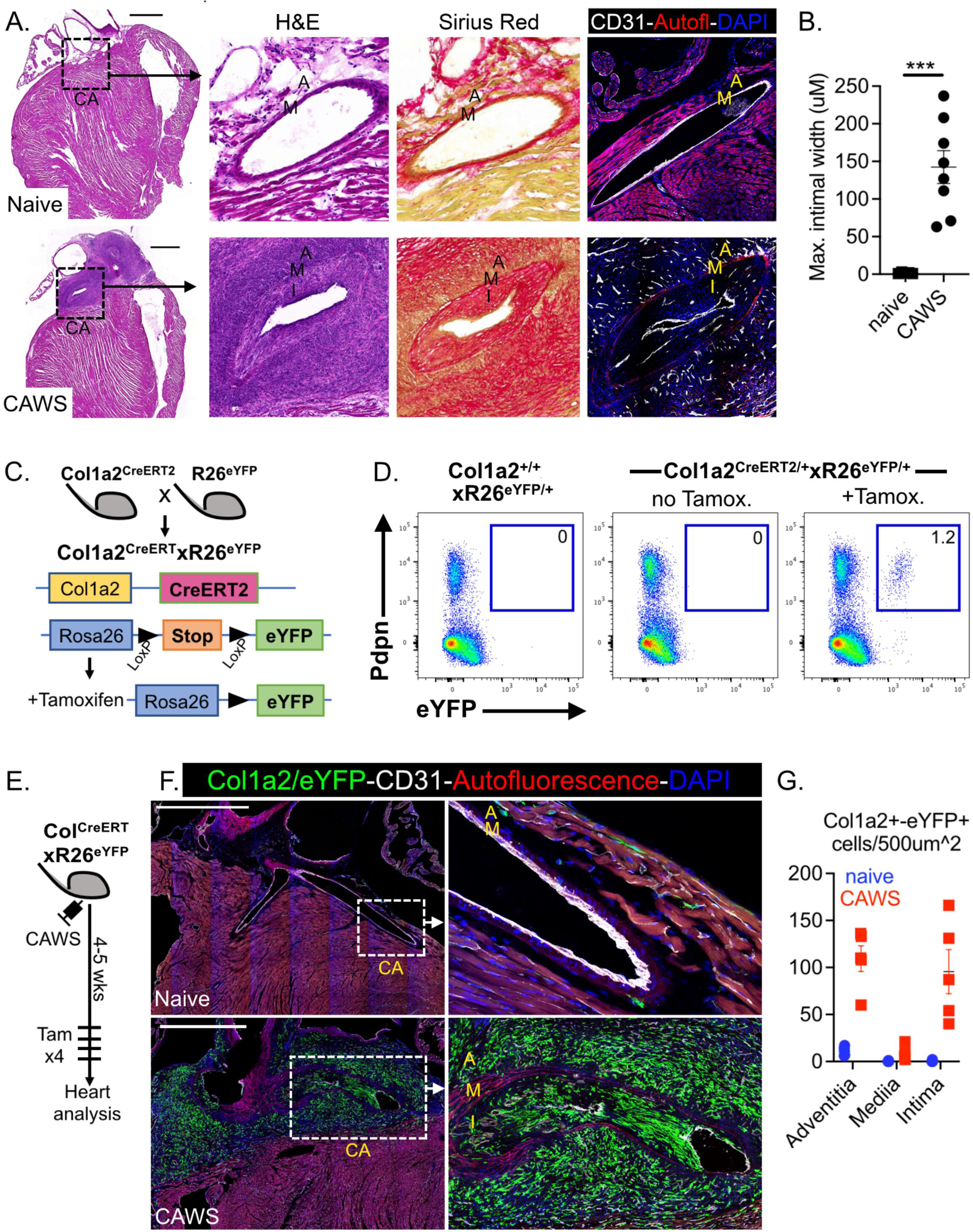
CAWS induced vasculitis triggers collagen-expressing fibroblasts to populate the coronary artery intima. **(A)** Cardiac sections from naïve or CAWS injected mice (4-5 weeks post injection) analysed by histology or immunofluorescent microscopy. For the latter, sections were stained for CD31 to label endothelial cells and autofluorescence was used to identify elastin fibres of the media. **(B)** Graphs show that maximum width of the coronary artery intima for individual mice (with mean±SEM) pooled from 3 independent experiments. **(C)** Genetic schema for Col1a2^CreERT2^.R26^eYFP^ system. **(D)** Flow cytometric analysis of cardiac cells from Col1a2^CreERT2^.R26^eYFP^ (and control) mice showing endogenous eYFP+ expression and anti-Podoplanin staining. Inset value is the mean of 4-6 mice from 3 independent experiments. **(E)** Experimental schema. **(F)** Cardiac sections from naïve or CAWS injected Col1a2^CreERT2^.R26^eYFP^ mice (4-5 weeks post injection) analysed by immunofluorescent microscopy. Sections were stained for GFP to identify Col1a2+/eYFP+ cells (green), CD31 to label endothelial cells (white) and autofluorescence to identify elastin fibres of the media (red). **(G)** Graphs show the number of Col1a2+/eYFP+ cells/500µm^2^ within each vessel layer for individual mice (with mean±SEM) pooled from 3 independent experiments. The coronary artery (CA), adventitia (A), media (M) and intima (I) are annotated and scale bars are 1000um.

We next determined if the intimal thickening observed in CAWS injected mice coincided with fibroblasts infiltration. To definitively identify fibroblasts, we used the Col1a2^CreERT2^ line, where the CreERT2 fusion gene is expressed under the control of a Collagen-1a2 (Col1a2) enhancer-promoter sequence [27]. To report Cre expression, Col1a2^CreERT2^ mice were crossed to the R26-stop-eYFP mice to create Col1a2^CreERT2^ x R26^eYFP^ mice. In this system, CreERT2 will undergo nuclear translocation upon tamoxifen administration and excise the LoxP flanked stop sequence, resulting in constitutive eYFP expression by Col1a2 expressing cells (**Fig 1C**). We first validated this system by flow cytometry, showing that eYFP+ cells emerged within the hearts of tamoxifen treated Col1a2^CreERT2/+^ x R26^eYFP/+^ mice (but not Cre negative controls or Col1a2^CreERT2/+^ x R26^eYFP/+^ mice without tamoxifen treatment) which expressed the fibroblast-marker Podoplanin (**Fig 1D**).

To determine if collagen-expressing fibroblasts infiltrate the coronary artery intima during CAWS-induced vasculitis, Col1a2^CreERT2^ x R26^eYFP^ mice were injected with CAWS and left for 4-5 weeks to allow intimal hyperplasia to develop. Tamoxifen was then administered to label Col1a2+ expressing fibroblasts for analysis by microscopy (**Fig 1E**). Cardiac sections were stained for eYFP to identify Col1a2+ fibroblasts, CD31 to identify endothelial cells and autofluorescence was used to identify the elastic fibres of the media. We found that the hearts of naïve mice contained a small population of widely distributed, Col1a2+ fibroblasts. These quiescent fibroblasts were detected within the myocardium and adventitial layer of the coronary arteries but were not found within the intimal layer (**Fig 1F-G**). By comparison, the hearts of CAWS injected mice contained a substantially expanded Col1a2+ fibroblast population which reproducibly localised to the aortic root and inflamed coronary arteries of CAWS mice. Critically, large numbers of Col1a2+ fibroblasts where present within both adventitial and intimal layers of the coronary arteries of CAWS mice (**Fig 1F-G**). These findings show that collagen-expressing fibroblasts expand and infiltrate the coronary artery intima during CAWS induced vasculitis, causing intimal hyperplasia.

### Intimal fibroblasts do not develop from epicardial-derived, resident fibroblasts

It has been suggested that intimal fibroblasts develop from their adventitial counterparts [16–18] and indeed, our findings that the emergence of intimal fibroblasts coincides with adventitial fibroblast expansion is in keeping with this possibility. We therefore used lineage tracing systems to determine if adventitial fibroblasts infiltrate the intimal layer during CAWS-induced vasculitis. To this end, we utilised the Wt.1^CreERT2^ line, which has CreERT2 inserted into the Wilms Tumour 1 (Wt.1) locus [28]. Wt.1 is a transcription factor that is temporally restricted to the epicardium during embryogenesis, and drives the epithelial-mesenchymal transition (EMT) of this population [29–31]. As such, this system allows the labelling of the epicardium and epicardial-derived cells (EPDCs), which others have shown includes resident cardiac fibroblasts [32]. To use this system, we crossed the Wt.1^CreERT2^ line to R26^eYFP^ mice and administered tamoxifen at E10.5 of pregnancy (**Fig 2A-B**). This schedule results in the selective labelling of the epicardium and their cardiac fibroblast progeny. To verify this approach, we confirmed that an eYFP+ cardiac population develops in E10.5 tamoxifen treated Wt.1^CreERT2/+^xR26^eYFP/+^ mice, but not Cre negative littermates (**Fig 2C**). Phenotypic analysis confirmed that the eYFP+ population had a fibroblastic phenotype, expressing high levels of podoplanin and PDGFRα, while being negative for CD31 and CD45 (**Fig EV1**). Moreover, RT-PCR analysis on sorted cardiac populations revealed that eYFP+ cardiac cells from Wt.1^CreERT2/+^xR26^eYFP/+^ mice express the highest levels of *Col1a2* mRNA (**Fig EV1**), confirming their identity as resident fibroblasts.

**Figure 2:**
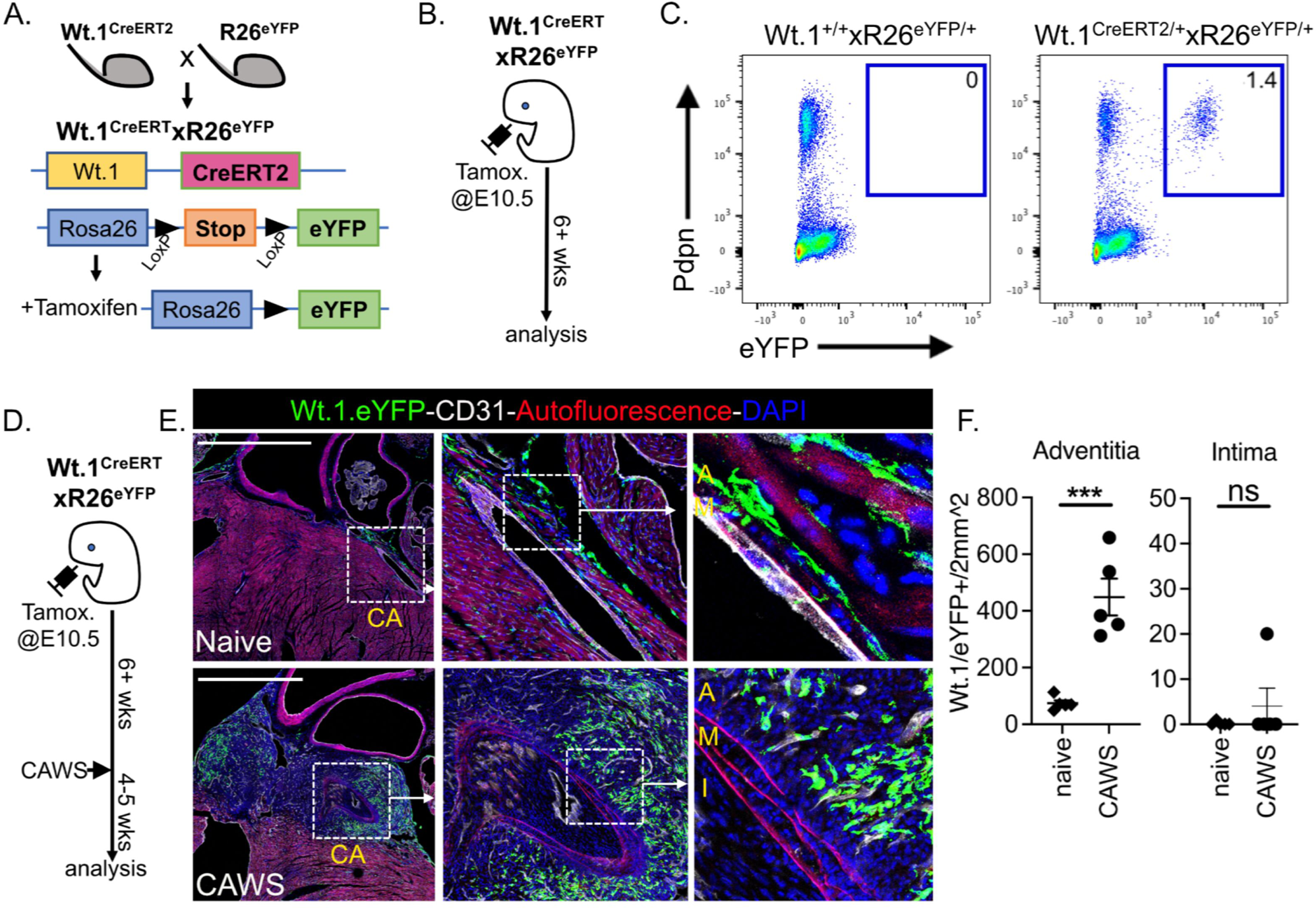
Epicardial-derived, resident fibroblasts do not infiltrate the coronary artery intima during CAWS induced vasculitis. **(A-B)** Genetic and experimental schema for Wt.1^CreERT2^.R26^eYFP^ system. **(C)** Flow cytometric analysis of cardiac cells from Wt.1^CreERT2^.R26^eYFP^ (and control) mice showing endogenous eYFP+ expression and anti-Podoplanin staining (mean of 5 mice/group from 3 independent experiments is inset). **(D)** Experimental schema. **(E)** Cardiac sections from naïve or CAWS injected Wt.1^CreERT2^.R26^eYFP^ mice (4-5 weeks post injection) analysed by immunofluorescent microscopy. Sections were stained for GFP to identify Wt.1+/eYFP+ cells (green), CD31 to label endothelial cells (white) and autofluorescence to identify elastin fibres of the media (red). **(F)** Graphs show the number of Wt.1+/eYFP+ cells/2mm^2^ within the coronary artery adventitia and intima for individual mice (with mean±SEM) pooled from 3 independent experiments. The coronary artery (CA), adventitia (A), media (M) and intima (I) are annotated and scale bars are 1000um.

We next analysed the distribution of Wt.1+/eYFP+ cardiac cells by confocal microscopy. This revealed that eYFP+ cells are widely distributed throughout the epicardial lining of the heart and the myocardium in the hearts of naïve Wt.1^CreERT2^xR26^eYFP^ mice (**Fig 2E**). Notably, Wt.1+/eYFP+ epicardial-derived cells (EPDCs) were enriched around the proximal coronary arteries which lie adjacent to the epicardium. These Wt.1+/eYFP+ EPDCs populate the adventitial layer of the coronary arteries, confirming robust labelling of adventitial fibroblasts using this system (**Fig 2E**).

To determine if epicardial-derived, adventitial fibroblast infiltrate the intima during CAWS-induced vasculitis, adult Wt.1^CreERT2^ x R26^EYFP^ mice (labelled at E10.5) were injected with CAWS and their hearts analysed by microscopy 4-5 weeks later (**Fig 2D**). This analysis revealed a significant increase in the eYFP+ population of CAWS injected Wt.1^CreERT2^x R26^EYFP^mice (**Fig 2E-F**). This population was highly concentrated in the adventitial layer around the coronary arteries of CAWS mice. However, despite adventitial fibroblast expansion, the intimal layer of the coronary artery was devoid of Wt.1+/eYFP+ EPDCs (**Fig 2E-F**). Thus, while epicardial-derived resident fibroblasts expand within the adventitial layer of the coronary arteries during vasculitis, this population does not migrate into the intima. These findings show that the intimal fibroblasts that emerge in the CAWS model of KD develop independently from adventitial fibroblasts.

### Intimal fibroblasts develop independently from endothelial cells

Endothelial cells have also been reported to give rise to myofibroblast in fibrotic disorders by undergoing endothelial-to-mesenchymal transition (endo-MT) [22, 23]. We therefore investigated if endothelial cells form intimal fibroblasts in CAWS-injected mice. To trace endothelial cells, we crossed VECad^CreERT2^ mice (where CreERT2 is driven by the vascular endothelial cadherin promoter) to the R26^eYFP^ reporter line, to achieve tamoxifen inducible eYFP labelling of endothelial cells (**Fig 3A**). As expected, the analysis of tamoxifen treated naïve VECad^CreERT2^xR26^eYFP^ mice by confocal microscopy showed extensive labelling of coronary artery CD31+ endothelial cells in the steady state (**Fig 3C**). To determine if this population undergoes endo-MT to generate intimal fibroblasts during vasculitis, tamoxifen treated VECad^CreERT2^xR26^eYFP^ mice were injected with CAWS and their hearts analysed by confocal microscopy 4-5 weeks later (**Fig 3B**). Despite profound intimal hyperplasia of the coronary arteries in CAWS injected mice, VECad+/eYFP+ cells remained localised to the luminal lining of the coronary artery, colocalising with the CD31+ endothelial cells (**Fig 3C**). Thus, endothelial cells did not infiltrate the intima during vasculitis, illustrating that intimal fibroblasts do not develop through endo-MT.

**Figure 3:**
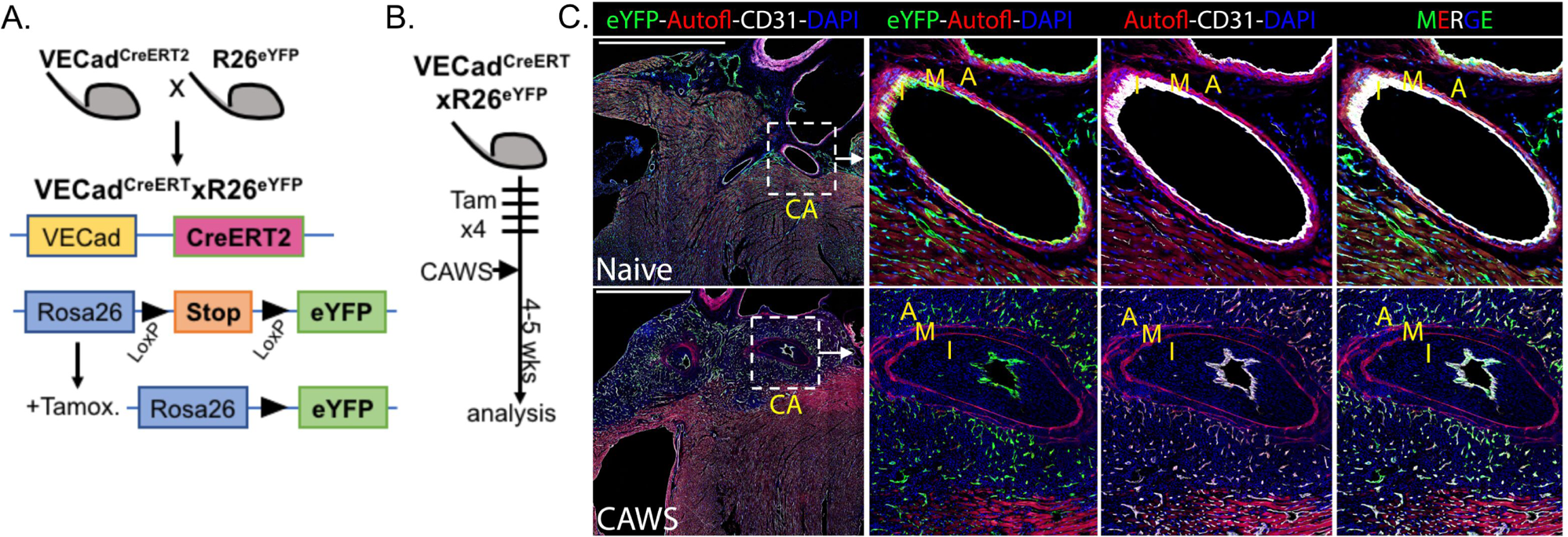
Endothelial cells do not form intimal fibroblasts during CAWS induced vasculitis. **(A-B)** Genetic and experimental schema for VECad^CreERT2^.R26^eYFP^ system. **(C)** Cardiac sections from naïve or CAWS injected VECad^CreERT2^.R26^eYFP^ mice (4-5 weeks post injection) analysed by immunofluorescent microscopy. Sections were stained for GFP to identify VECad+/eYFP+ cells (green), CD31 to label endothelial cells (white) and autofluorescence to identify elastin fibres of the media (red). Representative images of 5-6 mice per group (3 independent experiments) are shown with the coronary artery (CA), adventitia (A), media (M) and intima (I) are annotated and scale bars are 1000um.

### Intimal fibroblasts develop from Myh-11+ SMC precursors

Smooth muscle cells (SMCs) have been shown to form intimal fibroblasts during atherosclerosis [20], prompting us to examine whether a similar process occurs during vasculitis. To trace SMCs, we used Myh11^CreERT2^ mice [33], where the inducible CreERT2 is expressed under the control of the enhancer-promoter region from the smooth muscle myosin polypeptide 11 (*Myh11*), a contractile protein highly expressed by SMCs. To achieve tamoxifen inducible eYFP labelling of Myh11+ SMCs, Myh11^CreERT2^ mice were crossed to the R26^eYFP^ line (**Fig 4A**). Flow cytometric analysis confirmed that an eYFP+ population emerged within the hearts of tamoxifen treated, Myh11^CreERT2^xR26^eYFP^ mice but not Cre negative controls (**Fig 4B-C**). Phenotypic analysis revealed that the Myh11+/eYFP+ cardiac population was negative for phenotypic markers of endothelial cells (CD31), leukocytes (CD45) and fibroblasts (Pdpn, CD140a) but was uniformly positive for the mural cell marker CD146 (**Fig 4D**), confirming that Myh11^CreERT2^ recombination labels cardiac SMCs.

**Figure 4:**
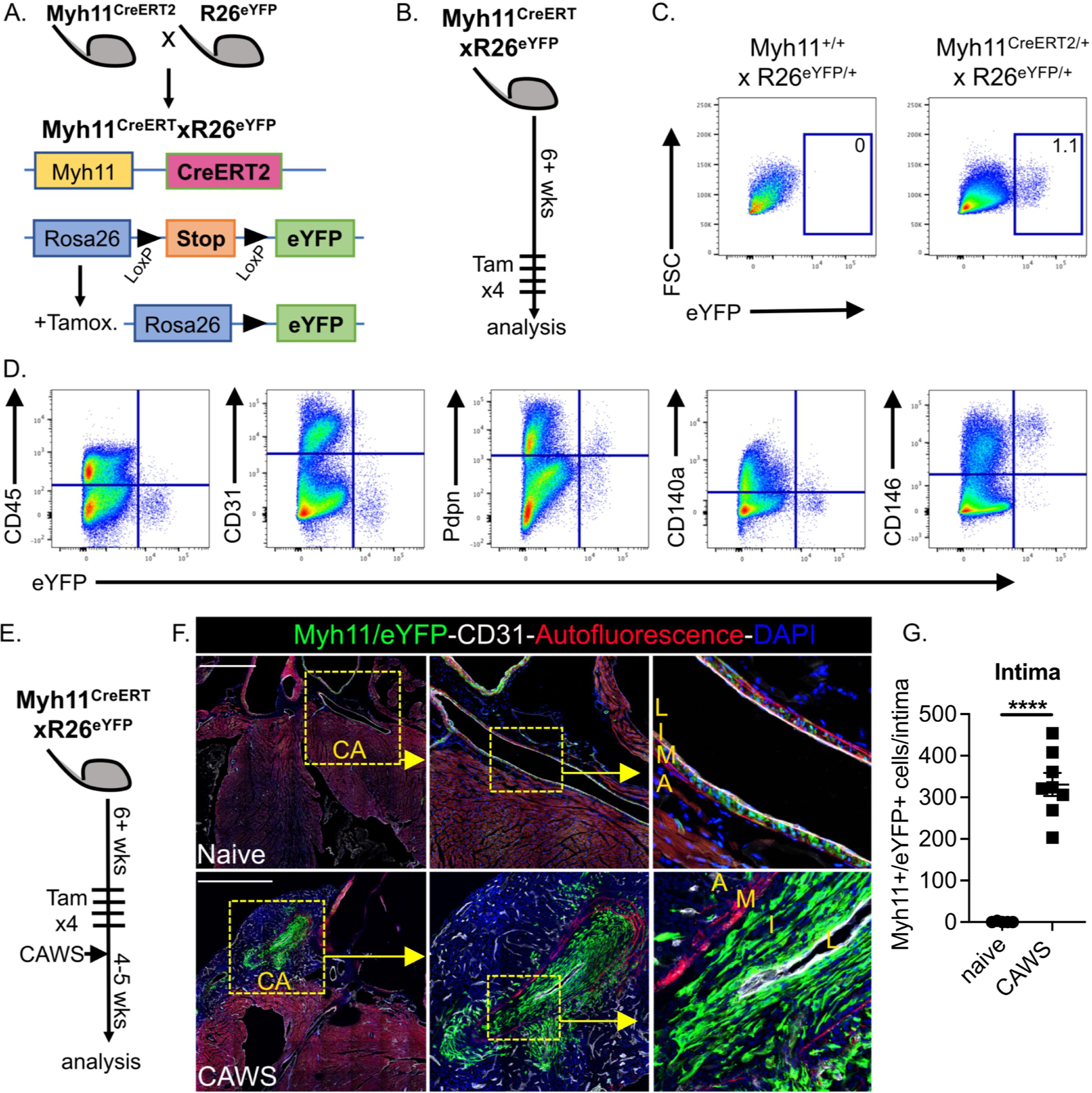
Myh11+ SMCs infiltrate the coronary artery intima during CAWS induced vasculitis. **(A-B)** Genetic and experimental schema for the Myh11^CreERT2^.R26^eYFP^ system. **(C-D)** Flow cytometric analysis of cardiac cells from Myh11^CreERT2^.R26^eYFP^ (and control) mice showing endogenous eYFP+ expression versus FSC (with the mean of 5-8 mice, 3 independent experiments inset**; (C)**) or a panel of lineage specific markers **(D)**. **(E)** Experimental schema. **(F)** Cardiac sections from naïve or CAWS injected Myh11^CreERT2^.R26^eYFP^ mice (4-5 weeks post injection) analysed by immunofluorescent microscopy. Sections were stained for GFP to identify Myh11+/eYFP+ cells (green), CD31 to label endothelial cells (white) and autofluorescence to identify elastin fibres of the media (red). **(G)** Graphs show the number of Myh11+/eYFP+ cells within the coronary artery intima for individual mice (with mean±SEM) pooled from 3 independent experiments. The coronary artery (CA), adventitia (A), media (M), intima (I) and lumen (L) are annotated and scale bars are 1000um.

To determine if SMCs form intimal fibroblasts during vasculitis, Myh11^CreERT2^ x R26^eYFP^ mice were administered tamoxifen to label SMCs and injected with CAWS. The hearts were analysed 4-5 weeks later by confocal microscopy (**Fig 4E**). As expected, Myh11+/eYFP+ cells were found exclusively within the medial layer of the coronary artery arteries in naive mice (**Fig 4F**). In contrast, Myh11-derived eYFP+ cells extensively populated the inflamed coronary artery intima of CAWS injected mice, forming the dominant population within the thickened intima (**Fig 4F**). The enumeration of confocal microscopy showed that Myh11-derived eYFP+ cells were significantly increased in the coronary artery intima of CAWS injected mice (**Fig 4G)**. Moreover, transcriptional analysis of eYFP+ cells sorted from the hearts of Myh11^CreERT2^ x R26^eYFP^ mice revealed that the Myh11-derived eYFP+ population had significantly increased expression of fibrillar collagens (*Col1a1, 1a2, Col3a1*) during CAWS induced vasculitis **(Fig EV2)**. These findings illustrate that vasculitis induces SMCs to undergo media-to-intimal migration and acquire a pro-fibrotic transcriptional program, resulting in the formation of intimal fibroblasts.

### The formation of SMC-derived intimal fibroblasts coincides with activation of the mTOR signalling pathway

We next investigated which signalling pathways control SMC-to-intimal fibroblast differentiation. We, and others, have reported that the mechanistic target of rapamycin (mTOR) pathway becomes activated by multiple populations within the inflamed arteries during vasculitis [34–37]. This prompted us to examine whether the SMC-derived intimal fibroblasts emerging during CAWS-induced-vasculitis utilise this signalling pathway. Cardiac sections from Myh11^CreERT2^ x R26^eYFP^ mice were stained for phosphorylated ribosomal protein S6 (pS6), a downstream target of (and biomarker for) active mTOR signalling [38]. We analysed cardiac sections from naïve controls and CAWS mice 21 days after injection, a timepoint when intimal fibroblasts are first emerging (**Fig 5A**). By staining for pS6, eYFP and Ki67 (to identify proliferating cells), we found that in naïve mice, Myh11+/eYFP+ cells were restricted to the media and negative for pS6 and Ki67, indicating minimal mTOR signalling or proliferation by quiescent SMCs (**Fig 5B**). By comparison, at day 21 post CAWS injection, Myh11+/eYFP+ cells had begun to infiltrate the intimal layer and activate mTOR signalling, as shown by strong pS6 staining. Notably, while pS6 staining was most striking in eYFP+ cells located adjacent to the endothelial layer (**Fig 5B**), we found that a majority of eYFP+ cells in both the media and intima were pS6+ of CAWS mice (**Fig 5C**). Thus, Myh11-derived cells in both layers of the vessel activate mTOR signalling during vasculitis. Finally, we found that a proportion of pS6+eYFP+ cells were Ki67+, linking mTOR activation with SMC proliferation. These findings reveal that the migration and expansion of SMC-derived intimal fibroblasts coincides with activation of the mTOR signalling pathway.

**Figure 5:**
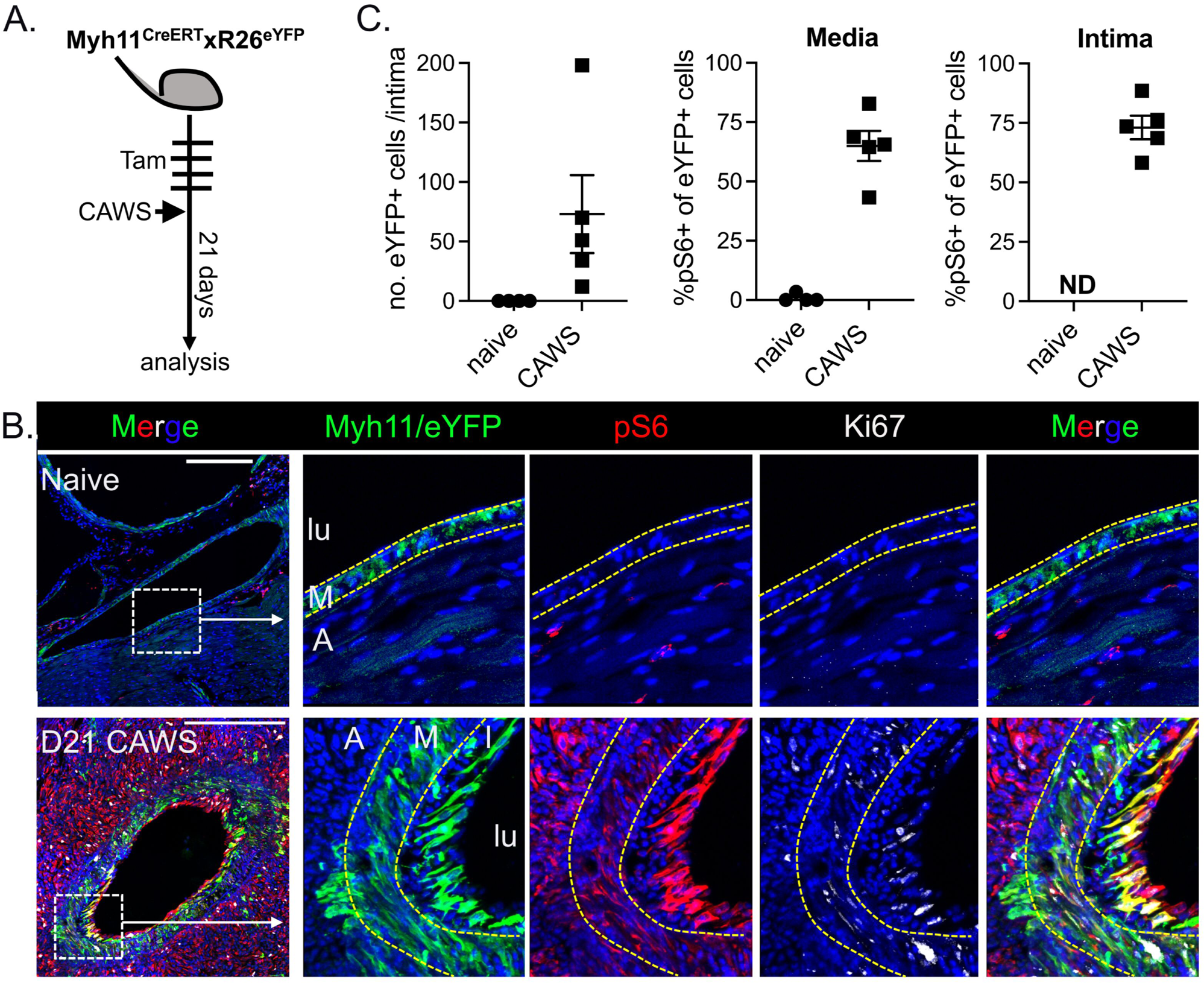
Myh11+ SMCs activate the mTOR signalling pathway during CAWS induced vasculitis drives. **(A)** Experimental schema. **(B)** Cardiac sections from naïve or day 21 CAWS injected Myh11^CreERT2^.R26^eYFP^ mice analysed by immunofluorescent microscopy. Sections were stained for GFP to identify Myh11+/eYFP+ cells (green), pS6 (red) to assess mTOR signalling and Ki67 (white) to measure cell proliferation. **(C)** Graphs show the number of Myh11+/eYFP+ cells within the coronary artery intima or % of Myh11/eYFP+ cells that are pS6+ within the media and intimal layers of the coronary artery. Dots represent individual mice (with mean±SEM) pooled from 2 independent experiments. The coronary artery (CA), adventitia (A), media (M), intima (I) and lumen (L) are annotated and scale bars are 250um.

### Intimal fibroblasts activate mTOR signalling in patients with Kawasaki disease, giant cell arteritis and Takayasu’s arteritis

We next examined if the mTOR pathway is also activated in the intimal fibroblasts that emerge in vasculitis patients. We first analysed coronary artery sections from autopsy tissue of two infants who died from myocardial infarction during acute KD. H&E staining revealed that in comparison to controls obtained from cardiac disease-free organ donors, the coronary arteries from both KD cases showed extensive immune cell infiltrate, which coincided with regions of intimal hyperplasia (**Fig 6A**). To identify mTOR signalling by intimal fibroblasts, coronary artery sections were stained for α−SMA (to identify myofibroblasts) and pS6 (to measure mTOR activity) and analysed by confocal microscopy. Control arteries from organ donors showed minimal pS6 staining by the α−SMA+ within the media and intima, indicating limited mTOR signalling in the steady state (**Fig 6B**). In comparison, both KD cases showed strong regional expression of pS6+ by α−SMA+ fibroblasts within the inflamed intima (**Fig 6B**). This analysis reveals that the mTOR signalling pathway is activated by the fibroblasts that populate the intimal layer of inflamed coronary arteries in acute KD.

**Figure 6:**
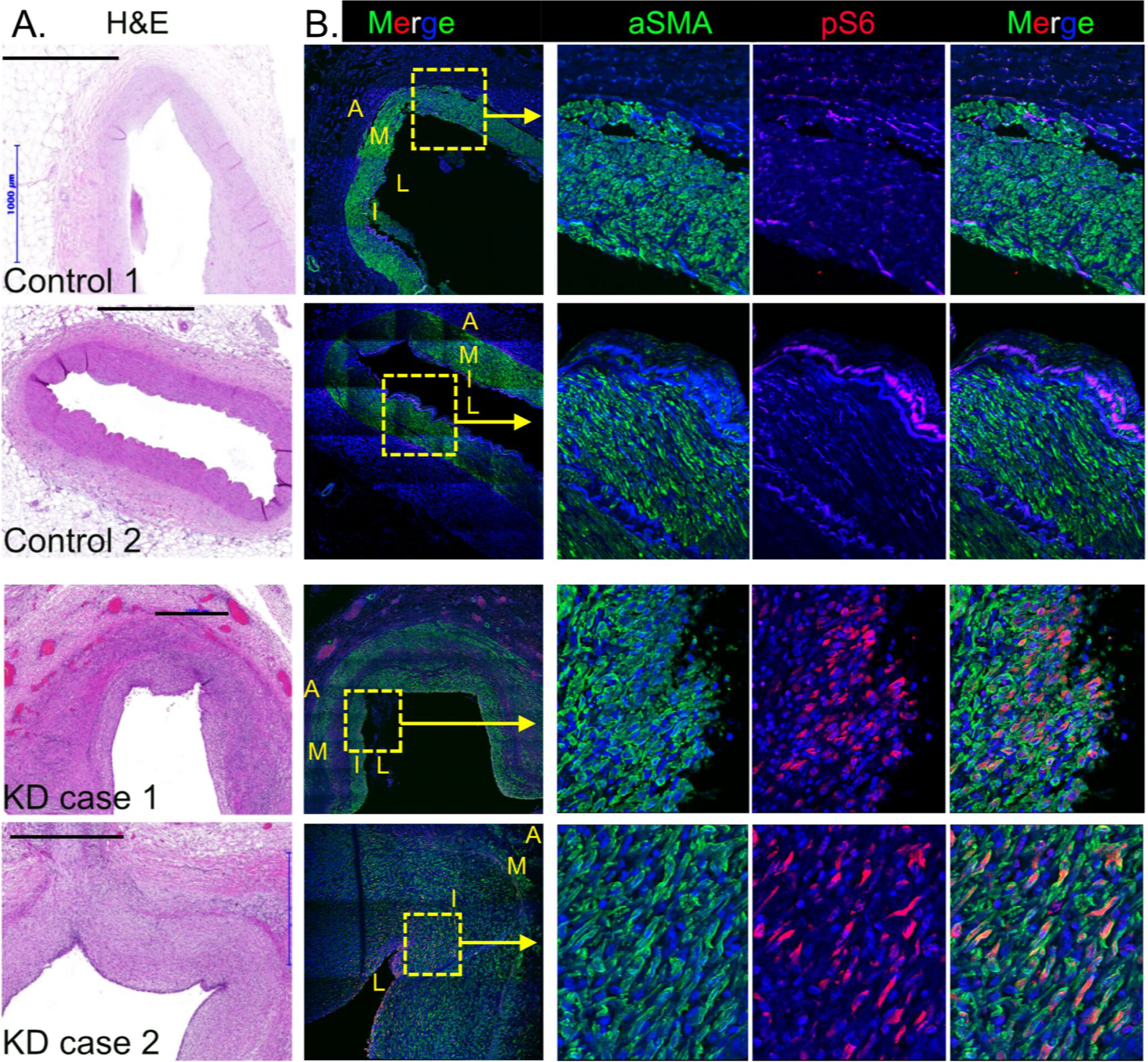
Intimal fibroblasts of KD patients have an activated mTOR signalling. Coronary artery sections from two cardiac-disease free organ donors or two acute KD fatalities analysed by H&E staining **(A)** or immunofluorescent microscopy **(B)**. For immunofluorescence, sections were stained for *a*-SMA (green) and pS6 (red) and analysed by confocal microscopy. Box shows inset area and the adventitia (A), media (M), intima (I) and lumen (L) are annotated and scale bars are 1000um.

We next investigated whether the pathogenic intimal fibroblasts that emerge in other forms of vasculitis also activate the mTOR signalling pathway. We first examined Takayasu’s arteritis (TAK), a large vessel vasculitis involving the aorta and its major branches [8]. H&E analysis of coronary artery sections from two TAK fatalities revealed profound remodelling of the coronary arteries, associated with intimal hyperplasia and immune cell infiltrate (**Fig 7A**). Probing for mTOR signalling by confocal microscopy revealed that pS6+/α-SMA+ intimal myofibroblasts were clearly evident in one of the TAK cases (TAK case 1; **Fig 7B**). We postulate that this region of pS6+ myofibroblasts corresponds to an area of nascent vascular remodelling, and that mTOR signalling is a feature of intimal fibroblast activation in TAK.

**Figure 7:**
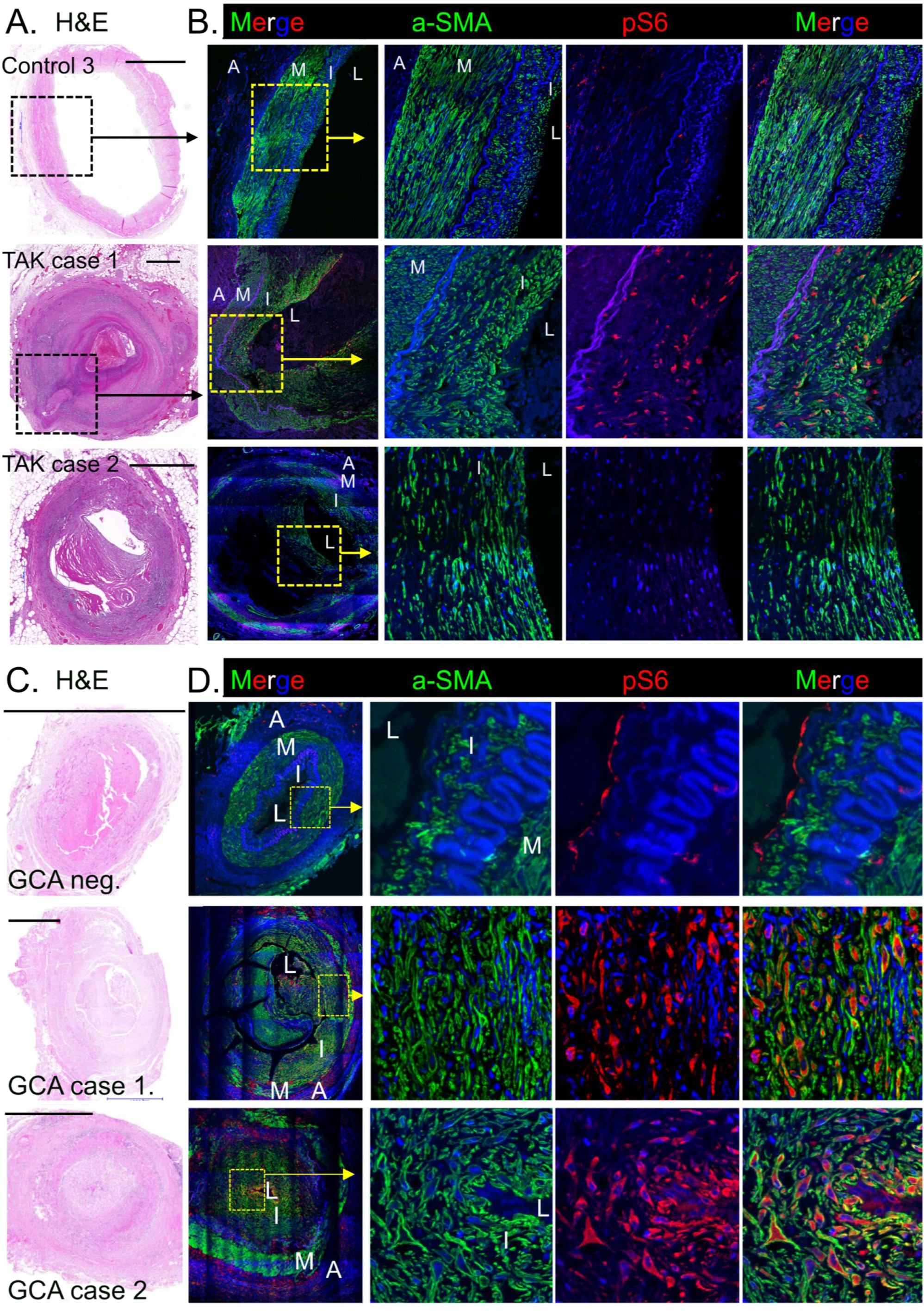
Intimal fibroblasts of patients with Takayasu’s arteritis and Giant Cell arteritis have an activated mTOR signalling pathway. **(A-B)** Coronary artery sections from one cardiac-disease free organ donor (control 3) and two Takayasu’s arteritis fatalities analysed by H&E staining **(A)** or immunofluorescent microscopy **(B)**. For immunofluorescence, sections were stained for *a*-SMA (green) and pS6 (red) and analysed by confocal microscopy. **(C-D)** Temporal artery sections from a GCA negative control and two GCA positive patients analysed by H&E staining **(C)** or immunofluorescent microscopy **(D)** as above. Box shows inset area and the adventitia (A), media (M), intima (I) and lumen (L) are annotated and scale bars are 1000um.

We also examined temporal artery sections from patients with giant cell arteritis (GCA). This systemic vasculitis occurs in the elderly and can affect the aorta and its major branches, most notably the carotid arteries and branching temporal arteries [8, 9]. We screened temporal artery sections from diagnostic biopsies, analysing two GCA positive cases together with a GCA negative, control temporal artery. H&E analysis revealed that the GCA positive arteries showed extensive intimal thickening and variable immune cell infiltrate (**Fig 7C**). Confocal analysis revealed that while α−SMA+ cells within the normal temporal artery were pS6 negative, a large proportion of α−SMA+ intimal myofibroblasts within the inflamed temporal arties of both GCA cases showed strong pS6 staining (**Fig 7D**). Collectively, these findings illustrate that the mTOR signalling is activated by the intimal fibroblasts that drive adverse vascular remodelling in patients with KD, TAK and GCA.

### mTORC1 signalling is an intrinsic and essential regulator of SMC-derived intimal fibroblast formation

To formally address whether mTOR signalling directly controls the development of intimal fibroblasts during vasculitis, we asked whether removing mTOR signalling from SMCs abrogated their ability to form intimal fibroblasts? To disrupt mTOR signalling, we genetically targeted the Regulatory Associated Protein of mTOR (Raptor), which is a major subunit of the mTORC1 complex [38–40]. To this end, we utilised Raptor-floxed mice, which have exon 6 of the raptor locus flanked by LoxP sites, allowing Cre-mediated *raptor* deletion [40]. To drive SMC-specific *raptor* deletion, Raptor^fl/fl^ mice were crossed to the Myh11^CreERT2^ line. Notably, the *raptor*^fl/fl^ line were also bred to carry the R26-stop-eYFP, meaning that Myh11-driven recombination will simultaneously delete *raptor* and activate eYFP transcription (allowing the identification of *raptor*-deficient SMCs through eYFP expression; **Fig 8A**). To investigate how SMC-specific raptor deletion impacts intimal fibroblasts formation, Myh11^CreERT2^xRaptor^fl/fl^xR26^eYFP^ mice (where Raptor is deleted from eYFP+/SMCs) and Myh11^CreERT2^xRaptor^+/+^xR26^eYFP^ controls (where Raptor is intact in eYFP+/SMCs) were administered tamoxifen to trigger SMC recombination and then injected with CAWs to induce vasculitis (**Fig 8B**). After 4-5 weeks, hearts were analysed by histology and confocal microscopy. H&E staining revealed that CAWS injection induced comparable levels of cardiac inflammation in both Raptor^+/+^ and Raptor^fl/fl^ groups (as measured by the cardiac area with immune cell infiltrate; **Fig 8C/F**). As in previous experiments, confocal microscopy showed that in control Myh11^CreERT2^xRaptor^+/+^xR26^eYFP^ mice (where Raptor was intact), Myh11+/eYFP+ cells robustly infiltrated the coronary artery intima following CAWS injection causing profound intimal hyperplasia (**Fig 8D-E**). In stark contrast, the Myh11+/eYFP+ cells of Myh11^CreERT2^ xRaptor^fl/fl^xR26^eYFP^ mice (where raptor was deleted in SMCs) were absent from the coronary artery intima and remained entirely within the medial layer (**Fig 8D-E**). Thus, despite robust cardiac inflammation, deleting raptor from SMCs completely abrogated their ability to infiltrate the coronary artery intima during vasculitis. Moreover, by measuring the size of the coronary artery intima, we found that SMC-raptor deficient mice had a significantly reduced intimal area following CAWS injection (compared to Raptor+/+ controls), demonstrating that inhibiting SMC migration reduces vasculitis-induced intimal hyperplasia (**Fig 8G**). These findings illustrate that (i) mTORC1 signalling is an intrinsic and essential process for the development of SMC-derived intimal fibroblasts, (ii) preventing the formation of SMC-derived intimal fibroblasts via targeting mTOR, significantly reduces arterial stenosis in vasculitis and (iii) mTORC1 therefore represents a rational therapeutic target in vasculitis.

**Figure 8:**
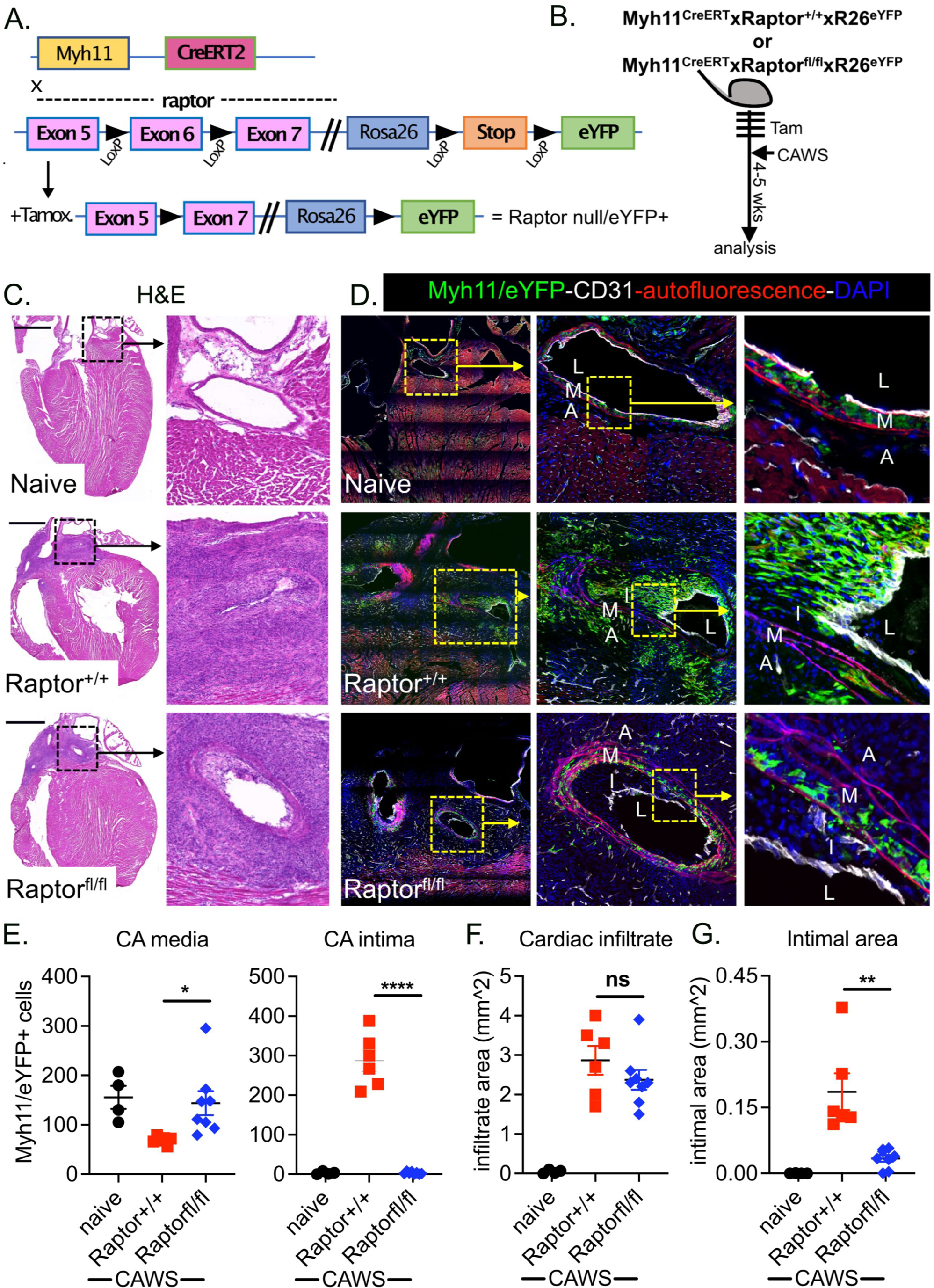
mTORC1 is an intrinsic and essential regulator of SMC-derived intimal fibroblast formation. **(A-B)** Genetic and experimental schema for Myh11^CreERT2^.Raptor^fl/fl^.R26^eYFP^ system. **(C-D)** Cardiac sections from naïve or CAWS injected Myh11^CreERT2^.Raptor^fl/fl^.R26^eYFP^ or Myh11^CreERT2^.Raptor^+/+^.R26^eYFP^ mice (4-5 weeks post injection) analysed by H&E staining **(C)** or immunofluorescent microscopy **(D)**. For immunofluorescence, sections were stained for GFP to identify Myh11+/eYFP+ cells (green), CD31 to label endothelial cells (white) and autofluorescence to identify elastin fibres of the media (red). **(E-G)** Graphs show the number of Myh11+/eYFP+ cells within the coronary artery media or intima **(E)**, the total cardiac area with immune cell infiltrate **(F)** or the area of the CA intima **(G)**. Points depict individual mice (with mean±SEM) pooled from 3 independent experiments. The coronary artery (CA), adventitia (A), media (M), intima (I) and lumen (L) are annotated and scale bars are 1000um.

## Discussion

We report two major findings on the functional biology of intimal vascular fibroblasts in vasculitis. Using an array of complimentary lineage tracing systems in a mouse model of KD, we provide definitive evidence that vasculitis-induced intimal fibroblasts develop from migratory SMCs. Moreover, we exclude both epicardial-derived resident fibroblasts and endothelial cells as potential sources. Our findings show that neither adventitial-to-intimal migration (by fibroblasts) nor endothelial-to-mesenchymal transition (endo-MT) occurs during vasculitis. Instead, intimal fibroblasts emerge through the media-to-intimal migration of SMCs, which proliferate and upregulate collagen expression to form a pathogenic fibroblast population within the inflamed intima.

Our study also shows that the formation of SMC-derived intimal fibroblasts is driven by activation of the mTOR signalling pathway. mTOR is a protein kinase that can form two distinct multimeric signalling complexes, mTORC1 and mTORC2, each composed of an mTOR subunit and various binding partners [38]. The mTORC1 complex includes raptor and promotes cell growth and proliferation by amplifying protein translation and lipid biogenesis through the phosphorylation of downstream targets such as S6K1 (p70-S6 Kinase 1) and 4E-BP1 (the eukaryotic initiation factor 4E binding protein 1) [38]. Here, we show that the mTOR signalling cascade is activated by intimal fibroblasts that associate with arterial remodelling in patients with Kawasaki Disease, Takayasu’s arteritis and giant cell arteritis. Moreover, when mTORC1 complex was selectively deleted in SMCs (through the Myh11^CreERT2^-driven deletion of raptor), raptor-deficient SMCs completely failed to populate the inflamed intima in the CAWS model of KD. These findings demonstrate that signalling through the mTORC1 complex is: (i) activated in multiple forms of human vasculitis and (ii) plays an intrinsic and essential role in driving the development of pathogenic SMC-derived intimal fibroblasts.

Our findings raise a number of mechanistic questions and present clinical opportunities. Foremost is how does mTOR control the formation of SMC-derived intimal fibroblasts? Our finding that raptor-deficient SMCs remain within the media despite vascular inflammation, indicates that mTOR signalling most likely controls the migration potential of SMCs. Previous studies have reported that rapamycin inhibits SMC migration *in vitro* [41], and our findings corroborate this *in vivo*. However, precisely how SMCs undergo media-to-intimal migration during vasculitis, and how this process is regulated by mTOR requires further investigation.

It is noteworthy that SMC-derived fibroblasts strongly activate mTOR signalling (as measured by pS6 staining) within the intima of both CAWS mice and vasculitis patients. The persistence of mTOR signalling after intimal invasion implies that mTOR may also regulate other pathological features beyond SMC migration. Cellular proliferation is a distinct possibility because (i) fibroblast expand within the intima of KD patients (and CAWS mice) [6] and (ii) mTORC1 is known to promote cell-cycle progression through (amongst others) increasing the expression of cell cycle factors such as cyclins [38]. Thus, while further mechanistic studies are required, mTOR probably plays a multifaceted role in the development of SMC-derived intimal fibroblasts, controlling the migration and proliferative potential of this pathogenic population.

Our findings also raise the question of which factors trigger mTOR activation during vasculitis? IL-1 has recently been reported to drive SMC differentiation and activation in a mouse model of KD [42]. Given that IL-1 can directly activate mTOR signalling [43], it is possible that an IL-1-mTOR axis triggers the formation SMC-derived intimal fibroblasts. In addition, Notch signalling has also been reported to drive mTOR activation by fibroblasts cells in rheumatoid arthritis [44], and T cells in vasculitis [37]. Intriguingly, in these instances, the Notch activating ligand Jagged-1, was reported to be expressed on activated endothelial cells [37, 44]. Given our observations that the intimal fibroblast adjacent to the endothelial cells show high mTOR activity (as seen by increased pS6 staining), activated endothelial cells may drive intimal fibroblast activation/formation through a Notch-mTOR signalling axis. Identifying upstream regulators of mTOR remains an important objective to identify novel therapeutic options that limit intimal fibroblast activation.

We suggest our findings have significant clinical implications. The emergence of proliferating fibroblasts within the intimal layer of inflamed arteries is a major concern for vasculitis patients, as cellular accumulation can result in luminal narrowing and arterial occlusion [6]. These events frequently occur in severe vasculitis. For instance, a majority of KD patients with giant coronary aneurysms develop arterial stenosis [4], creating a significant risk for myocardial infarction [3]. Therefore, treatments that prevent adverse vascular remodelling in such high-risk patients are required. Our findings that mTOR signalling is essential for intimal fibroblast formation illustrate that this pathway is the ideal therapeutic target to reduce arterial stenosis. Moreover, our findings that mTOR is activated in the vascular intimal fibroblasts of patients with KD, GCA and TAK, indicate that mTOR inhibition may be effective in each of these types of systemic vasculitis, all of which have major unmet treatment needs.

Targeting mTOR is a feasible treatment strategy given the availability of safe and effective inhibitors. Indeed, we have recently reported that pharmacological inhibition of mTOR (using rapamycin) prevented adverse vascular remodelling in the CAWS model of KD [34]. Our current study indicates that the mode-of-action for rapamycin is (at least in part) through inhibiting SMC activation. Moreover, mTOR inhibitors are now widely used in drug-eluting stents to prevent re-stenosis [38, 45], illustrating their anti-stenotic potential in humans. Hence, we suggest that mTOR is an intrinsic, essential and druggable pathway which is activated in the intimal vascular fibroblasts that drive adverse remodelling in vasculitis. We believe that these findings provide compelling rationale for clinical trials of mTOR inhibitors as a novel therapeutic strategy in systemic vasculitis.

## Material & Methods

### Mice, tamoxifen administration and the CAWS model of Kawasaki Disease

C57BL/6, Col1a2^CreERT2^ [27], Wt.1^CreERT2^ [28], R26.eYFP [46], VeCAD^CreERT2^ [47], Myh11^CreERT2^ [33] and Raptor^flox^ mice [40] were bred at the Walter and Eliza Hall Institute of Medical Research (WEHI, Australia) under specific pathogen-free conditions. To induce the nuclear translocation of CreERT2, adult mice were administered 4mg tamoxifen (dissolved in peanut oil) by oral gavage daily, for four consecutive days. The *Candida albicans* water-soluble (CAWS) complex was prepared as previously described [25, 26, 48]. To induce a Kawasaki-like disease, adult mice were injected intraperitoneally with either 4mg of CAWS once or with 3mg CAWS on two consecutive days. All procedures were performed at WEHI and approved by the WEHI Animal Ethics Committee.

### Histology and immunofluorescent analysis

For imaging of mouse tissue, hearts were perfused with PBS and then fixed in 2% paraformaldehyde (4h on ice), dehydrated in 30% sucrose (12-18h at 4C) and embedded in OCT (Tissue-Tek). Hearts were sectioned (10μM) in the coronal plane and analyzed by histopathology or immunofluorescence microscopy. For histology, sections were stained with H&E or Sirius Red. To enumerate histology, cardiac sections were analyzed with an annotation tool in CaseCenter software (3DHISTECH, Hungary). For immunostaining, cardiac sections were hydrated in PBS, permeabilized with 0.1% Triton-X and non-specific staining blocked with serum (Jackson Immunoresearch), BSA and Protein block (Dako). Sections were stained with primary antibodies against GFP (ab290; Abcam), CD31 (MEC13.3; BD), phosphorylated-S6Ser240/244 Ribosomal Protein S6 (pS6; clone D68F8; Cell Signalling) and Ki67 (SolA15; Invitrogen) before detection with fluorochrome conjugated secondary antibodies (Invitrogen or Abcam). Slides were counterstained with DAPI, imaged on a Zeiss LSM-880 Confocal Microscope and analyzed with ImageJ software, including quantification.

For human studies, sections from fatal KD and TAK cases were obtained from the Victorian Institute of Forensic Medicine (VIFM), GCA biopsies were obtained from the Royal Melbourne Hospital and normal cardiac tissue was obtained from the Australian Donation and Transplantation Biobank (ADTB). Paraffin sections (7μM) were dewaxed and subjected to citrate antigen retrieval. Sections were blocked with serum/BSA, TrueBlack Lipofusin Autofluorescence Quencher (Biotium) and Protein block (Dako) before staining with anti-pS6 (detected with fluorochrome conjugated secondary antibodies) and α-SMA-AF488 (1A4; Invitrogen). All procedures were approved by the VIFM, RMH, ADTB and WEHI Human Research Ethics Committee.

### Flow cytometry of cardiac tissue

For flow cytometric analysis, murine hearts were digested in type I collagenase (1mg/ml; Worthington) with DNase I (10μg/ml; Sigma). Single cell suspensions were then stained with directly conjugated mAbs against CD45.2 (104), CD31 (390), podoplanin (8.1.1), PDGFRα/CD140A (APA5) and CD146 (ME-9F1) (BD Bioscience, eBioscience or Biolegend). Propidium iodide (100ng/ml) was added immediately prior to data acquisition on a Fortessa (BD) FACS machine and analyzed using Flowjo software.

### Real time quantitative PCR

For gene expression analysis of purified populations, cells were sorted on FACSAria II (BD) and RNA was extracted using the RNeasy Micro Kit (Qiagen) and cDNA synthesized with SuperScript III Reverse Transcriptase (Invitrogen) using oligo-dT primers (Promega). Quantitative real time PCR was performed with Fast Sybergren Master mix (In vitrogen) with customised primers for Hrpt (forward - CCCTCTGGTAGATTGTCGCTTA; Reverse – AGATGCTGTTACTGATAGGAAATTGA), Col1a1 (forward - GAGCGGAGAGTACTG GATCG; reverse - GCTTCTTTTCCTTGGGGTTC), Col1a2 (forward – CAGCGAAGAACTCATACAGC; reverse - GACACCCCTTCTACGTTGT), Col3a1 (forward – ACGTAAGCACTGGTGGACAG; reverse – GGAGGGCCATAGCTGAACTG). Target gene expression was normalized to Hrpt (ΔCT) and relative expression converted by the 2^−ΔCT^ method.

### Statistical analysis

Statistical analysis was performed with Prism 6.0 (GraphPad Software) using unpaired, two-tailed Student *t* tests. Statistical significance levels are expressed as *\* P<0.05; ** P<0.01; *** P<0.001; ****P<0.0001*.

## Acknowledgement

We thank Edan Azzopardi, Tom Kitson and Lauren Wilkins (WEHI, Melbourne, Australia) for outstanding technical assistance and Drs Christine Biben, Felicity Jackling, Jane Visvader (WEHI, Melbourne, Australia) and Axel Kallies (The Peter Doherty Institute, Melbourne, Australia) for the provision of mice. Biospecimens used in this project were provided by the Australian Donation and Transplantation Biobank (ADTB). We acknowledge the contribution of the Victorian Liver Transplant Unit, DonateLife Victoria and the ADTB to this research project. We gratefully acknowledge the generosity of the deceased organ donors and their families in providing valuable tissue samples.

## Sources of Funding

This work was supported by an Arthritis Rheumatology Australia Project Grant, the Reid Charitable Trusts and the Australian National Health and Medical Research Council (NHMRC) Program Grant 1113577 and Clinical Practitioner Fellowship 1154235 (to I.P.W.). This study was made possible through Victorian State Government Operational Infrastructure Support and the Australian Government National Health and Medical Research Council Independent Research Institute Infrastructure Support scheme. The ADTB was funded by the Australian Center for Transplant Excellence and Research.

## Conflict of Interest

IW’s lab has received funding from CSL and Med-Immune for research on cytokine antagonists. The remaining authors declare no conflict of interest relating to this research.

**Expanded View Figure 1: E10.5 Wt.1 recombination labels collagen expressing cardiac fibroblasts. (A)** Flow cytometric analysis of cardiac cells from Wt.1^CreERT2^.R26^eYFP^ mice (administered tamoxifen at E10.5) showing endogenous eYFP+ expression versus lineage specific markers. **(B)** Col1a2 mRNA expression (measured by RT-PCR) by cell subsets sorted from the hearts of adult Wt.1^CreERT2^.R26^eYFP^ mice (administered tamoxifen at E10.5). Graphs show the mean±SEM pooled from 3 independent experiment.

**Expanded View Figure 2: Myh11+ SMCs upregulate collagen expression during CAWS induced vasculitis. (A)** Flow cytometry sorting strategy of cardiac cells from Myh11^CreERT2^.R26^eYFP^ mice. **(B)** RT-PCR analysis of collagen expression by CD146+/eYFP+ cells isolated from the hearts of naïve and CAWS injected Myh11^CreERT2^.R26^eYFP^ mice. Data shows the mean±SEM from 2-3 pools of mice (2-4 mice/pool) acquired in 3 independent experiments.

## Notes

### Competing Interest Statement

The authors have declared no competing interest.

